## Supplementary Data for "Early recovery of proteasome activity in cells pulse-treated with proteasome inhibitors is independent of DDI2"

**Table S1. The sequence of DDI2 KO clones of HAP1 cells used in this work.**

| Horizon product # | CRISPR effect | gRNA sequence |
|---|---|---|
| HZGHC000182**c006** | 4bp deletion | GCTCGAAGTCGGCGTCGACC |
| HZGHC000182**c023** | 1bp insertion | GCTCGAAGTCGGCGTCGACC |
| HZGHC000396**c010** | 41bp deletion | AATAGCTATGGAAGAGGCTC |

**Table S2. Source of chemicals and reagents.**

| Reagent | Source |
|---|---|
| Bortezomib | LC Laboratories |
| Carfilzomib | LC Laboratories |
| CB-5083 | Cayman Chemicals |
| CHAPS | ThermoFisher |
| Cycloheximide | Sigma-Aldrich |
| Digitonin | GoldBio |
| PhosSTOP | Roche |
| Proteasome-Glo$^{TM}$ | Promega |
| CellTiter-Glo$^{TM}$ | Promega |
| RNAsinPlus | Promega |
| Suc-LLVY-amc | Bachem |
| Resazurin | Sigma |
| All others | VWR |

**Table S3. The sequence of siRNAs used in the work**.

| Target | Sequence | Vendor | Catalog # |
|---|---|---|---|
| DDI2 siRNA10 | GGACAUGCUUAAACGGCAC | Dharmacon | J-032713-10-0050 |
| DDI2 siRNA12 | CAAGAAAGGAUUCGUCUGU | Dharmacon | J-032713-12-0050 |
| Non-targeting pool | UGGUUUACAUGUCGACUAA<br>UGGUUUACAUGUUGUGUGA<br>UGGUUUACAUGUUUUCUGA<br>UGGUUUACAUGUUUUCCUA | Dharmacon | D-00110-10-20 |

**Table S4. Antibodies.**

| IgG | Type | Host | Dilution | Company | Catalog # |
|---|---|---|---|---|---|
| **Primary** | | | | | |
| TCF11/NRF1 D5B10 | mAb | rabbit | 1:500 | Cell Signaling | 8052S |
| GAPDH D4C6R | mAb | mouse | 1:1000 | Cell Signaling | 97166S |
| β-actin 8H10D10 | mAb | mouse | 1:1000 | Cell Signaling | 3700S |
| DDI2 | pAb | rabbit | 1:5000 | Bethyl | A304-629A |
| **Secondary** | | | | | |
| Anti-rabbit HRP-linked | | goat | 1:1000 | Cell Signaling | 7074S |
| Anti-mouse HRP-linked | | goat | 1:1000 | Cell Signaling | 7076P2 |
| Anti-rabbit-Alexa fluor 647 | | goat | 1:3500 | Invitrogen | A32733 |
| Anti-rabbit-Alexa Fluor 680 | | goat | 1:3500 | Invitrogen | A20176 |
| Anti-mouse-IRDye® 800CW | | goat | 1:3500 | Li-COR | 926-32210 |

**Table S5. qPCR primers**.

| Subunit | Gene name | Primers | Primer sequences |
|---|---|---|---|
| α6 | PSMA1 | forward | AGAGCTTGCAGCTCATCAGA |
| | | reverse | CAAGACGAGACACAGGCAGT |
| α2 | PSMA2 | forward | GCCCCGATTACAGAGTGC |
| | | reverse | TGGACGAACACCACCTGA |
| α7 | PSMA3 | forward | GGTGCGCAACTCTACATGATTG |
| | | reverse | TCCTCTGCCTCGTATCAGTTATTAA |
| α3 | PSMA4 | forward | CATTGGCTGGGATAAGCA |
| | | reverse | ATGCATGTGGCCTTCCAT |
| α4 | PSMA7 | forward | CTGTGCTTTGGATGACAACG |
| | | reverse | CGATGTAGCGGGTGATGTACT |
| β7 | PSMB4 | forward | TCTCGGCCAGATGGTGAT |
| | | reverse | CACATAACCGAGGAAGCT |
| β5 | PSMB5 | forward | GCTTGCCAACATGGTGTATC |
| | | reverse | ATCATAGGCCTGCTCCACTT |
| β2 | PSMB6 | forward | CCAAGGAAGAGTGTCTGCAA |
| | | reverse | TGCATCAGTACAGGGCATCT |
| β1 | PSMB7 | forward | ATTGACCTCTGCGTCATCAG |
| | | reverse | CTGTTTCTTCCAGCACCTCA |
| Rpt2 | PSMC1 | forward | GATGACCTCTCTGGTGCTGA |
| | | reverse | CCCTTTCAGGGATTGAGAAA |
| Rpt5 | PSMC3 | forward | CCAGAATCATGCAGATCCAC |
| | | reverse | GCCTTCCATGTAGTCCTCGT |
| Rpt3 | PSMC4 | forward | CCGCCAGAAGAGATTGATTT |
| | | reverse | ACAATGTAGCGGTTTTCACG |
| Rpn2 | PSMD1 | forward | GGGGACCTCTTCAATGTCAA |
| | | reverse | TAGGCAGAGCCTCATTTGCT |
| Rpn10 | PSMD4 | forward | AGGAGGAGGCCCGGC |
| | | reverse | TCACTTCTTGTCTTCC |
| Rpn7 | PSMD6 | forward | CAGTCAGCTGCTGGAATCAT |
| | | reverse | TGGTACTGCCAGTTCTTGCT |
| Rpn6 | PSMD11 | forward | GCTGCTCTGGAAACAATTCA |
| | | reverse | ACAGATGCACCAAATGAGGA |
| Rpn5 | PSMD12 | forward | TTGGTCCACACTTGTTGAGG |
| | | reverse | TGAGAGAAAGGCTTCGGACT |
| Rpn11 | PSMD14 | forward | TTGGATGGAAGGTTTGACAC |
| | | reverse | CCACATGTTCCTCCAAATGA |
| | PGK1 | forward | AAAGTCAGCCATGTGAGCACT |
| | | reverse | CCACCCCAGGAAGGACTTTA |

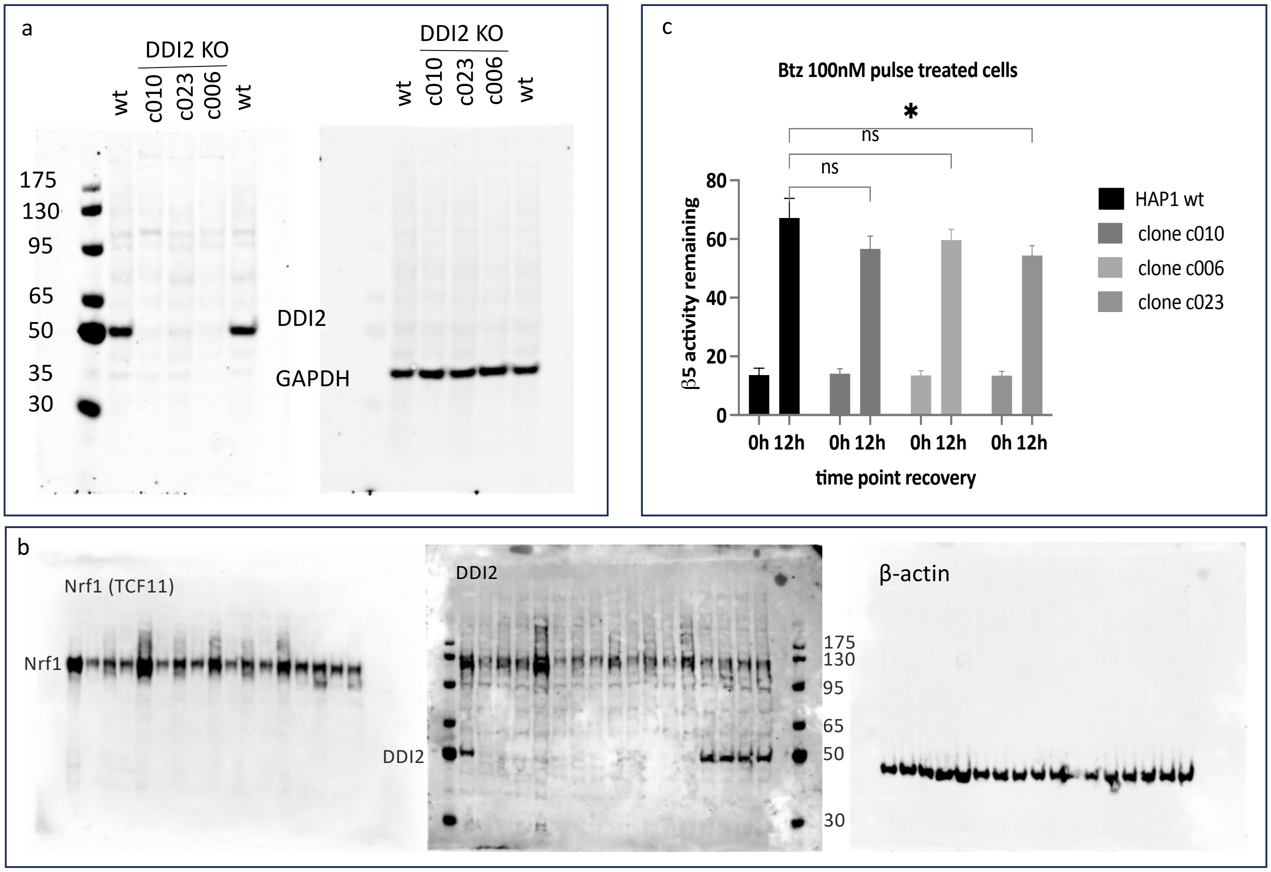


**Figure S1**, related to Figure 1. **a.** The full-size membrane for Fig. 1a. **b.** The full-size membranes for Fig. 1d. **c**. The proteasome activity of the samples used in Fig. 1d was measured with Suc-LLVY-AMC.

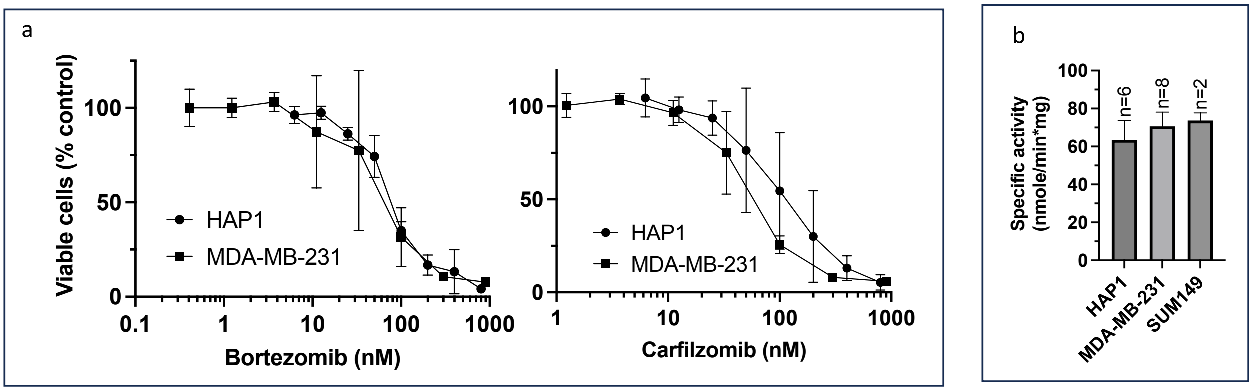


**Figure S2. Comparison of proteasome activity and PI sensitivity between HAP1, MDA-MB-231, and SUM 149 cells**. **a**. Cells were treated with PIs for 1h, media was shaken off, and cells were cultured in an inhibitor-free fresh media for 48h when Alamar Blue assay was performed; n=3-4. See Fig. 1 in [11] for a comparison of SUM149 and MDA-MB-231 cells. **b**. The β5 proteasome activity was measured using Suc-LLVY-AMC in the cell extracts of the untreated cell.

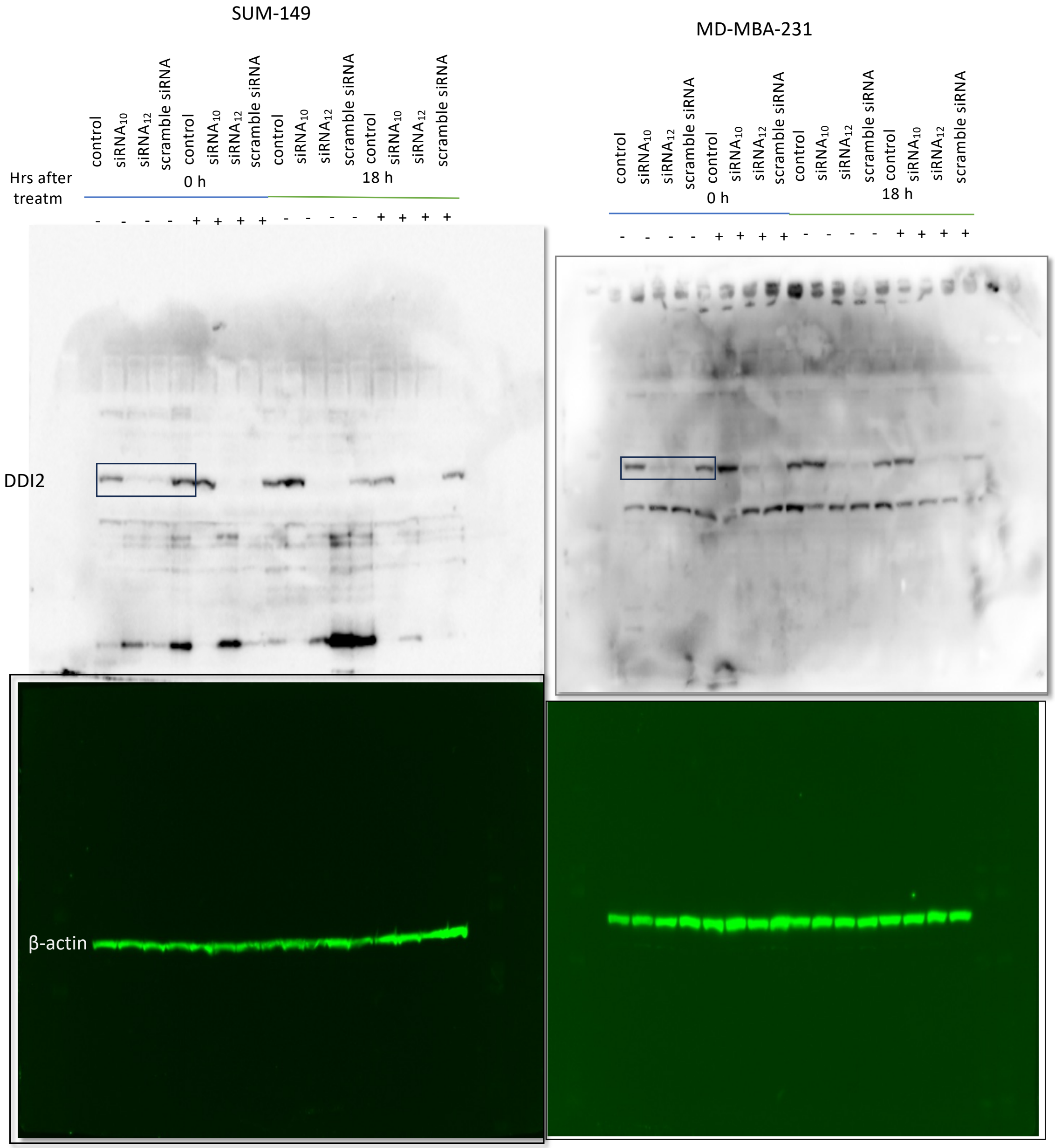


**Figure S3**, related to Figure 1. The full-size membranes of data on Fig. 1e. siRNA-transfected SUM-149 and MDA-MB-231 were treated with Btz for 1h. The cells were analyzed by western blot either immediately or 18h after 1hr Btz treatment. Boxes indicate data shown in Fig. 1e.

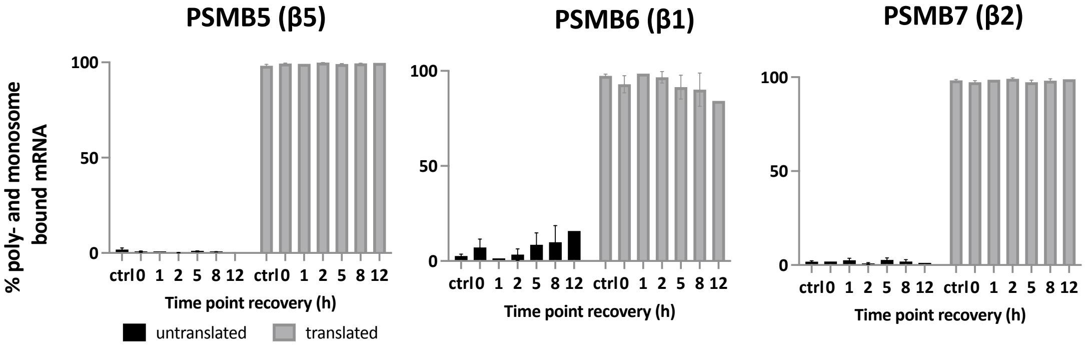


**Figure S4**, related to Fig. 3. Translation of catalytic subunits is not altered after treatment with inhibitors. Cells were treated with Btz for 1h, as in Fig. 3A, and then cultured in drug-free media. The cells were harvested at indicated times and analyzed by polysome profiling and qPCR, as shown in Fig. 3b; n=2.
